# Lineage Functional Types (LFTs): Characterizing functional diversity to enhance the representation of ecological behavior in Earth System Models

**DOI:** 10.1101/2020.01.01.891705

**Authors:** Daniel M. Griffith, Colin Osborne, Erika J. Edwards, Seton Bachle, David J. Beerling, William J. Bond, Timothy Gallaher, Brent R. Helliker, Caroline E.R. Lehmann, Lila Leatherman, Jesse B. Nippert, Stephanie Pau, Fan Qiu, William J. Riley, Melinda D. Smith, Caroline Strömberg, Lyla Taylor, Mark Ungerer, Christopher J. Still

**Affiliations:** Forest Ecosystems and Society, Oregon State University, OR, U.S.A.; US Geological Survey Western Geographic Science Center, Moffett Field, CA, 94035; NASA Ames Research Center, Moffett Field, CA, 94035; Department of Animal and Plant Sciences, University of Sheffield, U.K.; Department of Ecology and Evolutionary Biology, Brown University, RI, U.S.A.; Division of Biology, Kansas State University, KS, U.S.A.; South African Environmental Observation Network, National Research Foundation, Claremont, South Africa; Department of Biological Sciences, University of Cape Town, Rondebosch, South Africa; Department of Biology and the Burke Museum of Natural History and Culture, University of Washington, Seattle, WA, U.S.A.; Department of Biology, University of Pennsylvania, PA, U.S.A.; School of GeoSciences, University of Edinburgh, Edinburgh, U.K.; Department of Geography, Florida State University, FL, U.S.A.; Lawrence Berkeley National Laboratory, CA, U.S.A.; Department of Biology, Colorado State University, CO, U.S.A.

**Keywords:** C_4_ photosynthesis, earth system models, evolution, grass biogeography, plant functional types, vegetation models

## Abstract

Process-based vegetation models attempt to represent the wide range of trait variation in biomes by grouping ecologically similar species into plant functional types (PFTs). This approach has been successful in representing many aspects of plant physiology and biophysics, but struggles to capture biogeographic history and ecological dynamics that determine biome boundaries and plant distributions. Grass dominated ecosystems are broadly distributed across all vegetated continents and harbor large functional diversity, yet most Earth System Models (ESMs) summarize grasses into two generic PFTs based primarily on differences between temperate C_3_ grasses and (sub)tropical C_4_ grasses. Incorporation of species-level trait variation is an active area of research to enhance the ecological realism of PFTs, which form the basis for vegetation processes and dynamics in ESMs. Using reported measurements, we developed grass functional trait values (physiological, structural, biochemical, anatomical, phenological, and disturbance-related) of dominant lineages to improve ESM representations. Our method is fundamentally different from previous efforts, as it uses phylogenetic relatedness to create lineage-based functional types (LFTs), situated between species-level trait data and PFT-level abstractions, thus providing a realistic representation of functional diversity and opening the door to the development of new vegetation models.

## Introduction

Functional trait variation within and among biomes arises from environmental gradients and evolutionary histories that vary biogeographically, leading to species with differing ecological behavior and climatic responses, and differing impacts on ecosystem function, across continents (Lehmann *et al*., 2014; Higgins *et al*., 2016). Earth System Models (ESMs) typically apply abstracted plant functional types (PFTs; but see Scheiter *et al*., 2013; Pavlick *et al*., 2013; Medlyn *et al*., 2016) to represent physical, biological, and chemical processes crucial for soil and climate-related decision making and policy. However, PFTs must generalize across species, and inevitably encapsulate a wide range of plant strategies and vegetation dynamics, a demand that contrasts with efforts to investigate nuanced and species specific ecological behavior (Cramer *et al*., 2001; Bonan, 2008; Sitch *et al*., 2008; Kattge *et al*., 2011). Furthermore, PFTs account for only a modest degree of variation in a wide array of functional traits, ranging from seed mass to leaf lifespan (LL), in the TRY database (Kattge *et al*., 2011). For example, standard PFTs may not generally capture key drought responses in tree species (Anderegg, 2015), although models with a hydraulics module can be specifically tuned for this purpose (e.g., *ecosys;* Grant *et al*., 1995). Oversimplification of the physiognomic characteristics of PFTs can have major unintended consequences when simulating ecosystem function (Griffith et al., 2017), such as highly biodiverse savanna ecosystems (Searchinger *et al*., 2015). Studies that explicitly incorporate species-level trait variation into vegetation models (e.g., Grant *et al*., 1995; Lu *et al*., 2017) have demonstrated improvements in model performance. Selecting trait data from multi-variate trait distributions for model parameterization (Wang *et al*., 2012; Pappas *et al*., 2016) is very challenging for global modeling applications, particularly in hyper-diverse regions like the tropics, and may not be feasible for areas with biased or limited data. Until these data-gaps are filled, a finer-grained representation of the functional diversity among species might be achieved by reorganizing PFTs.

Importantly, in seeking approaches to restructure PFTs, numerous observations over the last decade have shown that both plant traits and biome-occupancy are commonly phylogenetically conserved, with closely related species having similar traits and niches (e.g., Cavender-Bares *et al*., 2009, 2016; Crisp *et al*., 2009; Liu *et al*., 2012; Donoghue & Edwards, 2014; Coelho de Souza *et al*., 2016). The existence of strong evolutionary constraints on plant functioning and distribution suggests that, as an alternative, vegetation types should be organized in a manner consistent with phylogeny. We advocate for explicit inclusion of evolutionary history and a consistent framework for integrating traits into global vegetation models. This approach brings a defensible method for defining vegetation types, enables the functional traits of uncharacterized species to be inferred from relatives, and allows evolutionary history to be explicitly considered in studies of biome history. Here, we illustrate this approach for grasses and grass-dominated ecosystems, where we use our framework to aggregate species into Lineage-based Functional Types (LFTs) to capture the species-level trait diversity in a tractable manner for large-scale vegetation process models used in ESMs. Capturing the evolutionary history of woody plants is also critical to understanding variation in ecosystems function in savannas (Lehmann *et al*., 2014; Osborne *et al*., 2018), and in general we are advocating for the development of LFTs in other vegetation types and in other ecosystems. Grasses provide a tractable demonstration for the utility of LFTs; we also discuss the potential to significantly improve ecological and biogeographical representations of other plants in ESMs.

Grasses are one of the most ecologically successful plant types on earth (Linder *et al*., 2018). Ecosystems containing or dominated by grasses (i.e., temperate, tropical, and subtropical grasslands and savannas) account for a large proportion of global land area and productivity. Grassy ecosystems cover some ~40% of earth’s ice-free surface (Gibson, 2009) and are a staple for humanity’s sustenance (Tilman *et al*., 2002; Still *et al*., 2003; Asner *et al*., 2004). The photosynthetic pathway composition (C_3_ or C_4_) of grass species is a fundamental aspect of grassland and savanna function, ecology, and biogeography. Of the ~11,000 grass species on earth, some ~4,500 use the C_4_ photosynthetic pathway (Osborne *et al*., 2014). Although they account for less than 2% of all vascular plant species (Kellogg, 2001), C_4_ grasses are estimated to cover some 19 million km^2^ and account for 20-25% of terrestrial productivity (Still *et al*., 2003), having risen to such prominence only in the last 8 million years (Edwards *et al*., 2010). Dominance by C_4_ versus C_3_ grasses has major influences on gross primary productivity and ecosystem structure and function (Still *et al*., 2003) and strongly influences interannual variability of the global carbon cycle, due to a combination of ecological and climatic factors (Poulter *et al*., 2014; Griffith *et al*., 2015). Dynamic vegetation models largely fail to reproduce spatial patterns of grass cover and productivity at regional to continental scales, limiting ability to predict future responses (Still *et al*., 2018). Just as often, these models also miss key transitions between biome states (Staver *et al*., 2011; Dexter *et al*., 2018; Still *et al*., 2018).

Most ESMs classify grasses into two PFTs based on differences between temperate C_3_ grasses and sub-tropical and tropical C_4_ grasses. However, grass ecological adaptations and physiological properties are highly diverse, ranging from cold-specialized to fire- and herbivore-dependent species. While grasses are often equated functionally, in reality they exhibit a high degree of variation in hydraulic, leaf economic, and phenological traits (Taylor *et al*., 2010; Liu *et al*., 2012) that likely explains their broad geographic dominance in different regions (Edwards *et al*., 2010; Visser *et al*., 2014). This includes economically important forest-forming grasses such as bamboos, but here we focus on globally dominant herbaceous lineages. Grasses exhibit strong phylogenetic diversity in leaf economics variation and associations with disturbance (Taylor *et al*., 2010; Liu *et al*., 2012; Simpson *et al*., 2016). Disturbances such as fire and herbivory have large impacts on ecosystem function and distributions, and PFT based approaches are unlikely to capture these differences among lineages. At broad phylogenetic and spatial scales, niche and biome conservatism of major plant lineages is common (Crisp *et al*., 2009; Cornwell *et al*., 2014; Donoghue & Edwards, 2014), and as such we argue that evolution and biogeography provide a framework for aggregating species (across ecosystems and strata) into LFTs that capture species-level trait diversity in a way that can be feasibly incorporated for use in global vegetation models, and that will improve PFT-based modeling approaches. Focusing on grasses, we developed this approach by collecting grass traits from databases (e.g., Osborne *et al*., 2011) and literature (e.g., Atkinson *et al*., 2016; Supplemental Appendix S1), for five key categories (physiology, structure, biochemistry, phenology, and disturbance). We summarize these species traits at the lineage level and relate these functional types to their observed global distributions.

## Methods for establishing lineage-based functional types (LFTs) for grasses

There are 26 monophyletic C_4_ lineages described in the Poaceae family, yet only two (the Andropogoneae and Chloridoideae) account for most of the abundance of C_4_ grasses globally (Lehmann *et al*., 2019; Fig 1.) (Edwards & Still, 2008; Edwards *et al*., 2010; Grass Phylogeny Working Group II, 2012). Among C_3_ grasses, only the Pooideae are globally dominant today. The Pooideae occupy cooler climates than the C_4_ Andropogoneae and Chloridoideae, which dominate in wetter and drier climates, respectively. Therefore, we focused on collecting species-level trait data from the literature and from databases for grass species from these three lineages. The term ‘trait’ is defined differently across research disciplines (Violle *et al*., 2007). Our aims necessitate a collection of broad trait space beyond that typically used for the leaf economic spectrum to include morphological and physiological determinants of plant hydraulics, physicochemical controls of photosynthesis, allocation to reproduction, and spectral reflectance. Many traits are highly connected, reflecting plant functional strategies. Further, a single trait can relate to multiple forms of plant fitness. Here, traits were assigned to groups in Table 1 based on their use in models and how they might be used in future applications (e.g., hyperspectral remote sensing of LFTs, or modeling of fire). We present median and variation in trait values among-species for three major grass lineages (LFTs) as per Figure 1, and compare these with commonly used values for C_3_ and C_4_ PFTs (Table 1).

**Figure 1.**
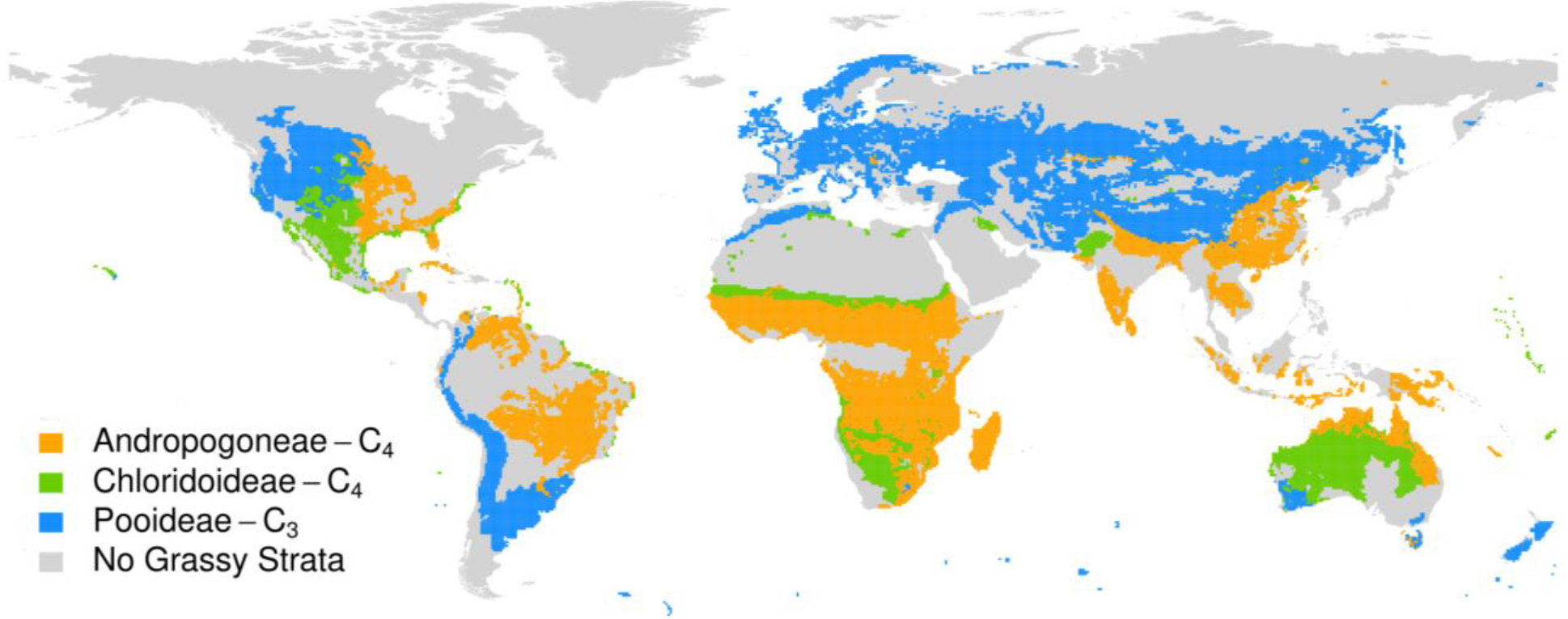
Distributions of the three globally dominant grass lineages in the herbaceous layer. These data come from Lehmann et al (2019).

**Table 1.**
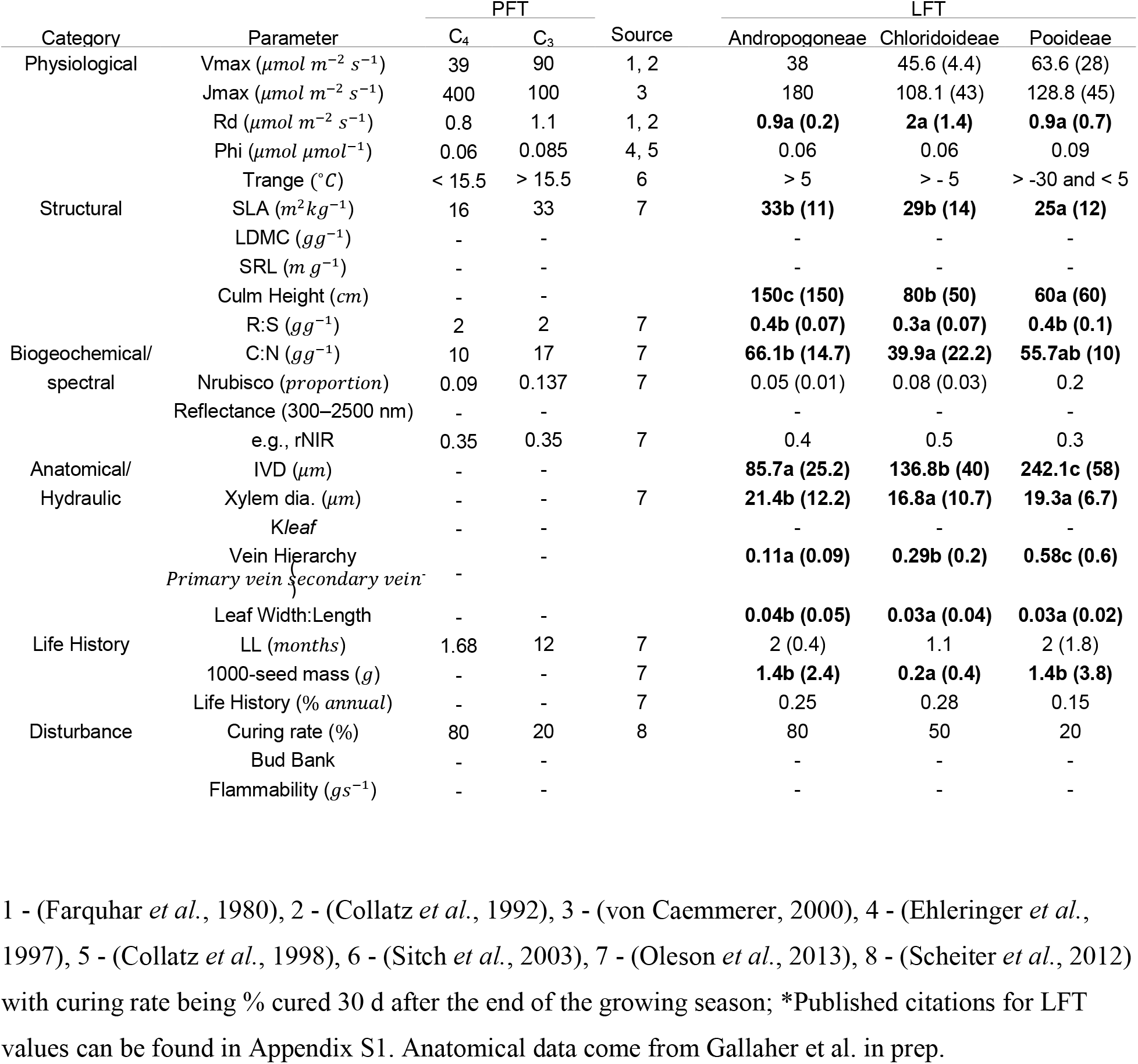
Common PFT parameters from ESM models, and median LFT parameters (IQR; interquartile range in parentheses) for three dominant grass lineages, taken from the literature and trait databases. Lineage assignments are based on Osborne *et al*. (2014). The table shows a subset of common parameters, with up to five parameters from each of six major categories. Blank parameters are not typically included in many models but are potentially important for accounting for the ecological behavior of grasses. Letters indicate significant differences with a Tukey’s test from simple linear model fits when all three lineages had at least three data points. Sources are in table footer.

## LFTs for grasses differ drastically in key functional traits

Our LFTs demonstrate both the importance of considering lineage to explain ecological patterning, and the need for modification of current ESM PFT approaches. For instance, C_4_ plants typically have lower RuBisCO activity (V_max_) but higher electron transport capacity (J_max_) than C_3_ plants, reflecting both the additional energetic cost of C_4_ physiology and the greater efficiency of RuBisCO in higher CO_2_ environments (Collatz *et al*., 1998). The Chloridoideae grasses have intermediate V_max_ and J_max_ compared to the Andropogoneae and the Pooideae. Furthermore, the Pooideae have evolved to tolerate much colder conditions (Sandve & Fjellheim, 2010; Vigeland *et al*., 2013; McKeown *et al*., 2016), and our results suggest that C_4_ lineages may differ in their thermal tolerances (Watcharamongkol *et al*., 2018). These differences suggest that macroecological synthesis studies with global implications (e.g., Walker *et al*., 2014; Heskel *et al*., 2016) should, at minimum, include more grass species in their datasets, ideally organized around LFTs.

Trade-offs among adaptations and tolerances in natural systems are believed to promote coexistence among plant species (Tilman, 1988; Tilman & Pacala, 1993; Kneitel & Chase, 2004). Specific leaf area (SLA) measures the cost of constructing a leaf, which represents a tradeoff between acquisitive (high relative growth rate) and conservative (high leaf lifespan) plant strategies (Westoby, 1998; Westoby *et al*., 2002; Wright *et al*., 2004). Model simulations of growth are highly dependent on the value of SLA used (Korner, 1991; Sitch et al., 2003; Bonan, 2008). However, in most ESMs, C_3_ grass PFTs have higher or similar SLA values as C_4_ PFTs likely biasing predictions. In contrast, we found that the C_4_ LFTs had higher SLA than the C_3_ LFT, but SLA did not differ between the two dominant C_4_ grass lineages (Atkinson et al. 2016). SLA can be highly variable within lineages in grasses, likely due to the importance of herbivore pressure as a competing demand on leaf economics (Anderson *et al*., 2011; Griffith *et al*., 2017) as well as intraspecific variation. As a result, SLA highlights that some traits are harder to generalize than others using the LFT approach, and suggests that a range of values may be appropriate than a single value for constraining LFT parameters. The phylogenetic signal among grass lineages is stronger for stature (Taylor *et al*., 2010; Liu *et al*., 2012), with the Andropogoneae being considerably taller on average than the Chloridoideae. This difference suggests that not all traits are oriented along a fast-slow axis at broad taxonomic scales across C_3_ and C_4_ grass lineages (Reich, 2014; Díaz et al., 2016). Furthermore, the C_3_- and eudicot-centric approach in the current leaf economics framework would suggest that a higher SLA should also correlate with a higher specific leaf nitrogen content, yet the evolution of C_4_ photosynthesis allows for a significant reduction in Rubisco content, and hence plant nitrogen requirements (Taylor 2010). Thus, grass lineages differ in numerous leaf traits which have consequences that extend from palatability and flammability to hydrological differences.

Physiological and morphological leaf vascular traits underlie variation in SLA, constrain the hydrology of plants (e.g., Blonder *et al*., 2014; Sack *et al*., 2014), and are key traits related to the evolution of C_4_ photosynthesis (Sage, 2004; Ueno, 2006). We show here key hydraulic differences between the two dominant C_4_ lineages, which correspond to the C_4_ biochemical subtypes (Ueno, 2006; Liu & Osborne, 2015). The Chloridoideae have low conductance and high embolism resistance hydraulic traits (Table 1), and tend to inhabit drier sites (Fig. 1). Some Andropogoneae have been described as “water spenders” (Williams *et al*., 1998), and their hydraulic traits help to explain their affinity with higher rainfall habitats where they rapidly expend available soil water (Taub, 2000) and promote fire after curing. These hydraulic differences should have large effects in models, especially those that consider tree-grass coexistence (Higgins *et al*., 2000) and explicit representation of plant hydraulics (Grant *et al*., 1995).

Lineages also differ in biogeochemical traits that influence nutrient turnover rates and the reflectance and absorbance properties of vegetation. For example, Andropogoneae have higher C:N than Chlordoideae grasses, a likely result of growth rate differences and the frequent association of Andropogoneae grasses with fire. Similarly, a greater proportion of N in Chloridoideae leaves is allocated to RuBisCO, which is related to V_max_ (Ghannoum et al. 2012). Finally, C_3_ and C_4_ grasses are distinguishable spectrally at the leaf, canopy, and landscape level based on differences between the functional types in chlorophyll a/b ratio, canopy structure, and seasonality (Foody & Dash, 2007; Siebke & Ball, 2009; Irisarri *et al*., 2009). C_3_ and C_4_ grasses are typically given many of the same optical properties in vegetation models, but we show here that Chloridoideae might have considerably higher NIR reflectance than other lineages, possibly producing interesting optical variation and affecting the surface energy balance (Ustin & Gamon, 2010)(Table 1). Foliar spectral traits are also correlated with morphological and chemical traits related to nutrient cycling and plant physiology (Dahlin *et al*., 2013; Serbin *et al*., 2014).

Grass lineages also show key differences in phenological and reproductive traits that should be captured in models, especially those that include demographic predictions (Davis *et al*., 2010). Chloridoideae grasses have seeds with lower mass than other lineages (Liu *et al*., 2012; Bergmann *et al*., 2017), and this may represent a trade-off with higher seed production and other ‘fast’ growth strategies (Adler *et al*., 2014). Wind versus animal dispersal strategies might also affect diaspore size in a way not directly related to disturbance, whereas some reproductive traits may also indicate fire and disturbance-related adaptations. Phenological traits exhibit conservatism across many plant lineages (Davies *et al*., 2013). Fire and herbivory are two globally important and contrasting disturbances for grass-dominated vegetation (Archibald & Hempson, 2016). It is less clear how herbivory effects can be captured in such models, given that these herbivore-related traits vary greatly in grasses (Anderson *et al*., 2011). Many fire-related traits show patterns of phylogenetic conservatism, with high flammability clustering into particular lineages such as the Andropogoneae (Simpson *et al*., 2016). Large-scale vegetation models that have simulated grass fires in Africa have attributed faster curing (becoming dry fuel) rates to C_4_ vegetation (Scheiter *et al*., 2012), and this behavior appears to be due largely to dominant Andropogoneae grasses.

We have identified large differences among LFTs, across six trait categories, that are not captured by the standard PFT approach. Many of these trait data have very low sample sizes (from 1 to 1365) and come from non-overlapping species, highlighting the need for systematic data collection for grasses. Such a data collection effort would be an excellent opportunity to test for coordination among trait axes in a phylogenetic context, which has rarely been done in other systems despite the likelihood that relatedness drives patterns of trait covariation (e.g., Salguero-Gómez *et al*., 2016; Griffith *et al*., 2016). Furthermore, intra-groups trait variation deserves to be properly estimated (only some traits in Table 1 have enough data to estimate variability) as convergence and adaptation produce meaningful trait variation that should be incorporated into models.

## Potential for lineage-based functional types in other vegetation types

Many current PFTs implicitly represent groupings of closely related lineages (e.g., pinaceous conifers, grasses). However, even in these cases biogeographic distributions, and the coarseness of the phylogenetic unit, generates a lack of useful resolution. Currently, there are efforts to incorporate species-level trait data and methods such as those proposed by Cornwell *et al*., (2014) could be employed to cluster species into prominent lineage-based groupings representing unique trait combinations. Phylogenies are hierarchical by nature and allow the LFT approach to be scalable and adjustable to the research question being addressed. The ability to remotely sense plant lineages adds potential for rapidly developing LFTs from spectral data (e.g., Cavender-Bares *et al*., 2016). LFTs would be valuable for a wide range of systems. For example, trees in Eurasian boreal forests have evolved to suppress canopy fires, whereas North American boreal trees enable greater intensity canopy fires. These distinctions lead to major differences in CO2 emissions and function (Rogers *et al*., 2015) that might be captured in an LFT framework. Secondly, LFTs for savanna tree communities could better represent differing climatic responses that are driven by unique evolutionary and biogeographic histories (Lehmann *et al*., 2014; Osborne *et al*., 2018). Finally, tropical ecosystems such as the dipterocarp forests in Southeast Asia would be well suited to LFTs which might better represent carbon storage (Brearley *et al*., 2016).

Potential challenges with a lineage-based functional approach include the fact that many plant traits do not show strong phylogenetic conservatism (Cadotte *et al*., 2017). There are likely spatial and phylogenetic scales at which the LFT approach will be most appropriate; for example, at large scales (regional to continental), lineage conservatism is common (Crisp *et al*., 2009). In contrast, at the scale of local communities, we might expect character displacement and limiting similarity (processes that lead to reduced trait similarity of coexisting species) could obscure phylogenetic patterns and limit the utility of LFTs as proposed here (Webb *et al*., 2002; Cavender-Bares *et al*., 2009; HilleRisLambers *et al*., 2012). However, in grassy ecosystems, there is evidence that the patterns of spatial ecological sorting of lineages would be captured with LFTs also at landscape scales (e.g., within Serengeti National Park, Anderson *et al*., 2011; Forrestel *et al*., 2017). Finally, we focus on extant lineages that are functional important today, but their past interactions with other clades may have shaped the biomes they inhabit.

## Conclusions

We conclude that LFTs better capture functional diversity than PFT groupings, especially at the spatial scales common for ESMs (e.g., 100 km). Our analysis of current knowledge of grass functional (physiology, structure, biochemistry, phenology, and disturbance) diversity, distributions, and phylogeny indicates that to represent grass ecological behavior in ESMs, at least two C_4_ and at least one C_3_ LFTs are required of today’s most ecologically dominant grasses. These proposed LFTs capture key evolutionary differences in physiological, structural, biogeochemical, anatomical, phenological, and disturbance-related traits. We also highlight the need for systematic trait data collection for grasses, which we show are vastly underrepresented in trait databases, despite their ecological and economic importance. More broadly, we outline the LFT framework which is highly flexible and has the potential for use in a wide range of applications. We advocate for the use of phylogeny as a way to help guide and constrain the inclusion of burgeoning plant trait data to expand the range of functional types considered by global vegetation models.

## Acknowledgements

This work was in part inspired by a workshop at the National Evolutionary Synthesis Center. CJS, DMG, SB, MU, FQ, JBN were supported by National Science Foundation award 1342703; CAES and TJG by NSF awards 1253713, 1342787, and 1120750. CERL was supported by an award from The Royal Society. WJR was supported as part of the RUBISCO Scientific Focus Area in the Regional and Global Climate Modeling Program of the U.S. Department of Energy, Office of Science, Office of Biological and Environmental Research under contract DE-AC2–5CH11231.

